# Role of C2-domain protein CAR1 in the plasma membrane ABA signaling response

**DOI:** 10.1101/2023.09.18.558310

**Authors:** Ai-Yu Guo, Wen-Qiang Wu, Di Bai, Yan Li, Jie Xie, Siyi Guo, Chun-Peng Song

**Affiliations:** State Key Laboratory of Crop Stress Adaptation and Improvement, School of Life Sciences, Henan University, Kaifeng 475004, China; Sanya Institute of Henan University, Sanya 572025, Hainan, China

**Author notes:** These authors contributed equally (Ai-Yu Guo, Wen-Qiang Wu). The author responsible for distribution of materials integral to the findings presented in this article in accordance with the policy described in the Instructions for Authors (https://academic.oup.com/plphys/pages/General-Instructions) is Chun-Peng Song. **One-sentence summary:** CAR1 recruits three major components of early ABA signaling to the plasma membrane using membrane microdomains as localization regions.

**Keywords:** abscisic acid, C2-domain ABA-related protein, Single-molecule, Membrane microdomains

## Abstract

Plasma membrane (PM)-associated abscisic acid (ABA) signal transduction is an important component of ABA signaling. The C2-domain ABA-related (CAR) proteins have been reported to play a crucial role in recruiting ABA receptor PYR1/PYL/RCAR (PYLs) to the PM. However, the molecular details of the involvement of CAR proteins in membrane-delimited ABA signal transduction remain unclear. Here, the GUS-tagged materials for all *Arabidopsis* CAR members were used to comprehensively visualize the extensive expression patterns of the *CAR* family genes. Based on the specificity of CAR1’s response to ABA, we determined to use it as a target to study the function of CAR proteins in PM-associated ABA signaling. Single-particle tracking showed that ABA affected the spatiotemporal dynamics of CAR1. The presence of ABA prolonged the dwell time of CAR1 on the membrane and showed faster lateral mobility. Surprisingly, we verified that CAR1 could directly recruit Hypersensitive to ABA1 (HAB1) and SNF1-related protein kinase 2.2 (SnRK2.2) to the PM at both the bulk and single-molecule levels. Furthermore, PM localization of CAR1 was demonstrated to be related to membrane microdomains. Collectively, our study revealed that CARs recruited the three main components of ABA signaling to the PM to respond positively to ABA. This study deepens our understanding of ABA signal transduction.

## Introduction

Abscisic acid (ABA) is a hormone that is typically associated with plant major stress response (Chen et al., 2020). The ABA signaling pathway involves three main components: ABA receptors (PYR/PYL/RCARs, hereafter referred to as PYLs), type 2C protein phosphatases (PP2Cs) (negative regulator), and SNF1-related protein kinase 2s (SnRK2s) (positive regulator). When ABA is present, PYLs bind with ABA to create a complex in which PP2Cs participate, allowing PP2Cs to release SnRK2s protein kinase inhibition. Subsequently, SnRK2s are activated either by themselves or by other protein kinases, such as Raf-like MAPKKKs (Nguyen et al., 2019). SnRK2s control a variety of physiological responses by phosphorylating their target substrates, including transcription factors, ion channels, and transporters (Wang et al., 2013). In the absence of ABA, PP2Cs interact with and repress SnRK2s, thereby blocking the ABA signaling pathway.

Numerous ABA signaling elements have been discovered and investigated. The ABA receptor family in *Arabidopsis* consists of 14 PYLs. PYLs selectively interact with PP2Cs and exhibit distinct binding characteristics with ABA (Tischer et al., 2017). Although PYLs have redundant functions in regulating ABA co-receptor PP2Cs, some PYLs have individual functions in regulating different downstream factors. The 80 PP2Cs in *Arabidopsis* include nine clade A PP2Cs and three clade E PP2Cs, which act as negative regulators of stress responses. In particular, under non-stress or moderate stress conditions, clade A PP2Cs are key negative regulators of ABA signaling that inhibit stress signaling and enable the proper level of plant growth inhibition (Komatsu et al., 2013). ABA-activated SnRK2s play key roles in regulating the transcriptional response to ABA, and knockout mutants of the three subtypes of SnRK2 members (*snrk2*.*2snrk2*.*3snrk2*.*6*) are extremely insensitive to ABA (Fujita et al., 2009). SnRK2s undergo autophosphorylation and self-activation after being released from PP2Cs, which then activate or inhibit a number of downstream transcription factors and PM proteins, including NADPH oxidase, anion channels SLAC1, SLAH1, and SLAH3, K^+^ transporter AKT1, and PM water channel protein (AQP) AtPIP2 (Chérel et al., 2002; Sirichandra et al., 2009; Geiger et al., 2011; Brandt et al., 2012; Grondin et al., 2015).

ABA can be detected both intracellularly and on the plasma membrane (PM) (Zhu, 2016). However, the core ABA receptor PYLs may not bind to the PM directly and may need assistance from other proteins to sense the ABA on the PM. In a recent study, the PYL4-interacting protein C2 domain ABA-related 1 (CAR1) was identified by a yeast two-hybrid (Y2H) screen using highly expressed PYL4 as bait in *Arabidopsis* (Rodriguez et al., 2014). PYL4 interacted with CAR1 in both the PM and nucleus of plant cells. And the membrane recruitment of PYLs by CAR1 occurred in a Ca^2+^-dependent manner. CAR1 is a member of a plant-specific gene family in *Arabidopsis* that encodes proteins ranging from CAR1 to CAR10. Structural and biochemical experiments have shown that CARs are peripheral membrane proteins that pierce the bilayer and functionally assemble on the membrane, providing a reasonable platform for ABA receptor membrane assembly (Diaz et al., 2016; Chen et al., 2023). In ABA-mediated related phenotypes, combinatorial mutants of CAR family members have reduced sensitivity to ABA compared to wild-type (WT), and there is functional redundancy between *CAR* genes, so triple mutants need to be generated to obtain robust phenotypes (Rodriguez et al., 2014).

The CAR protein family is crucial for regulating biotic and abiotic stress responses. Functional studies of CAR family members have been conducted in multiple species, including in rice (*Oryza sativa*), sweet potato (*Ipomoea batatas*), and *Arabidopsis*. CAR family members are involved in responses to salinity, wounding, blue light, and ABA, in addition to root gravitropic responses (Rodriguez et al., 2014; Dummer et al., 2016; Cheung et al., 2020; Dummer et al., 2021; You et al., 2022). In rice, GTPase activating protein 1 (OsGAP1) acts as a regulator of OsYchF1 to enhance biotic resistance (Cheung et al., 2013). In *Arabidopsis,* overexpression of the rice CAR homolog Small C2 domain protein 1 (OsSMCP1) improves salt and drought tolerance (Yokotani et al., 2009). In sweet potatoes, IbCAR1 contributes to increased salt tolerance by activating the ROS scavenging system through the ABA signaling pathway (You et al., 2022). In addition to CAR1 and CAR4, CAR9 has also been reported to be involved in the ABA response in *Arabidopsis*. The stability and localization of CAR9 are regulated by Lower temperature 1 (LOT1), thereby affecting ABA signaling to enhance plant tolerance to drought stress (Qin et al., 2019). CAR6, generally known as Enhanced bending 1 (EHB1), is the most studied member of CARs in *Arabidopsis*, although data on the involvement of EHB1 in ABA-related processes are still lacking. Ca^2+^-activated EHB1 acts as a central processor-like component that connects gravitropism and phototropism and works in concert with Nonphototropic hypocotyl 3 (NPH3) and ADP-ribosylation factor GTPase-activating protein 12 (AGD12) to achieve photogravitropic equilibrium (Knauer et al., 2011; Dummer et al., 2016; Michalski et al., 2017; Dummer et al., 2021). Genetic and physiological analyses have suggested that EHB1 acts as a negative regulator of iron acquisition by interacting with the iron transporter protein Iron-regulated transporter 1 (IRT1) (Khan et al., 2019).

Based on the current studies, CAR proteins can recruit ABA receptor PYLs to the PM. In addition, CAR proteins interact with some PM localization proteins to take part in appropriate biological processes. However, the mechanism of CARs in ABA signal transduction has not yet been fully elucidated. First, there are 10 CAR family genes in *Arabidopsis*; however, their tissue expression is unclear, which is fundamentally important for their functions. Second, the dynamics of CARs in response to exogenous ABA stimulation remain poorly understood. Third, it is unclear how ABA signaling is transduced from the PM. In addition to PYLs, whether CAR proteins can recruit other ABA signaling molecules (PP2Cs and SnRK2s) to the membrane, thereby directly responding to ABA signals on the membrane. Finally, the PM of cells contains multiple special regions, and it is not clear whether the binding of CAR to the PM is region-selective.

In this study, we first comprehensively revealed the tissue localization of CAR family genes using GUS-tagged materials. Furthermore, we used CAR1 as an example to study the dynamics of CAR proteins at the single-molecule level. Sequential images show that CAR1 responds positively to the presence of ABA. In addition, we confirmed that CAR1 could directly recruit HAB1 and SnRK2.2 to the PM. Finally, we showed that PM localization of CAR1 depends on the membrane microdomains. This study reveals the importance of CAR proteins in PM-associated ABA signaling.

## Results

### Spatial and temporal expression profiles indicate functional diversity among the *Arabidopsis CAR* family

Determining the expression of a gene in different tissues is crucial to investigating its function (Gonzalez-Guzman et al., 2012). Expression data available on the eFP Browser indicate that *CAR* family genes are expressed in different tissues (Winter et al., 2007). Using β-glucuronidase (GUS) as a reporter gene, we examined the transcription pattern of *CAR* family genes in *Arabidopsis* to verify the expression profiles of *CARs* (Figure 1). This analysis confirmed that *Arabidopsis CAR* family genes, represented by *CAR1*, *CAR3*, *CAR6*, *CAR8*, and *CAR10*, were expressed in different developmental stages and tissues (i.e., embryos, seedlings, primary roots, leaves, guard cells, flowers, and seed pods). *CAR5* and *CAR9* were only expressed in flowers, and specifically were highly expressed in pollen (Figure 1). Hypocotyl and root expression, especially vascular bundle expression, provided evidence of the involvement of CAR proteins in the responses to photogravitropic balance and ion transport (Khan et al., 2019; Dummer et al., 2021). In comparing the expression patterns of *CAR* family genes and ABA receptor *PYLs*, we found that the range of *CAR* gene expression covered the range of *PYL* gene expression (Gonzalez-Guzman et al., 2012). Therefore, CARs were involved in the signal perception of ABA on the PM at different developmental stages and in different tissues. Furthermore, they were widely expressed throughout the growth and development of *CAR* family genes, which suggests that they have powerful functions.

**Figure 1.**
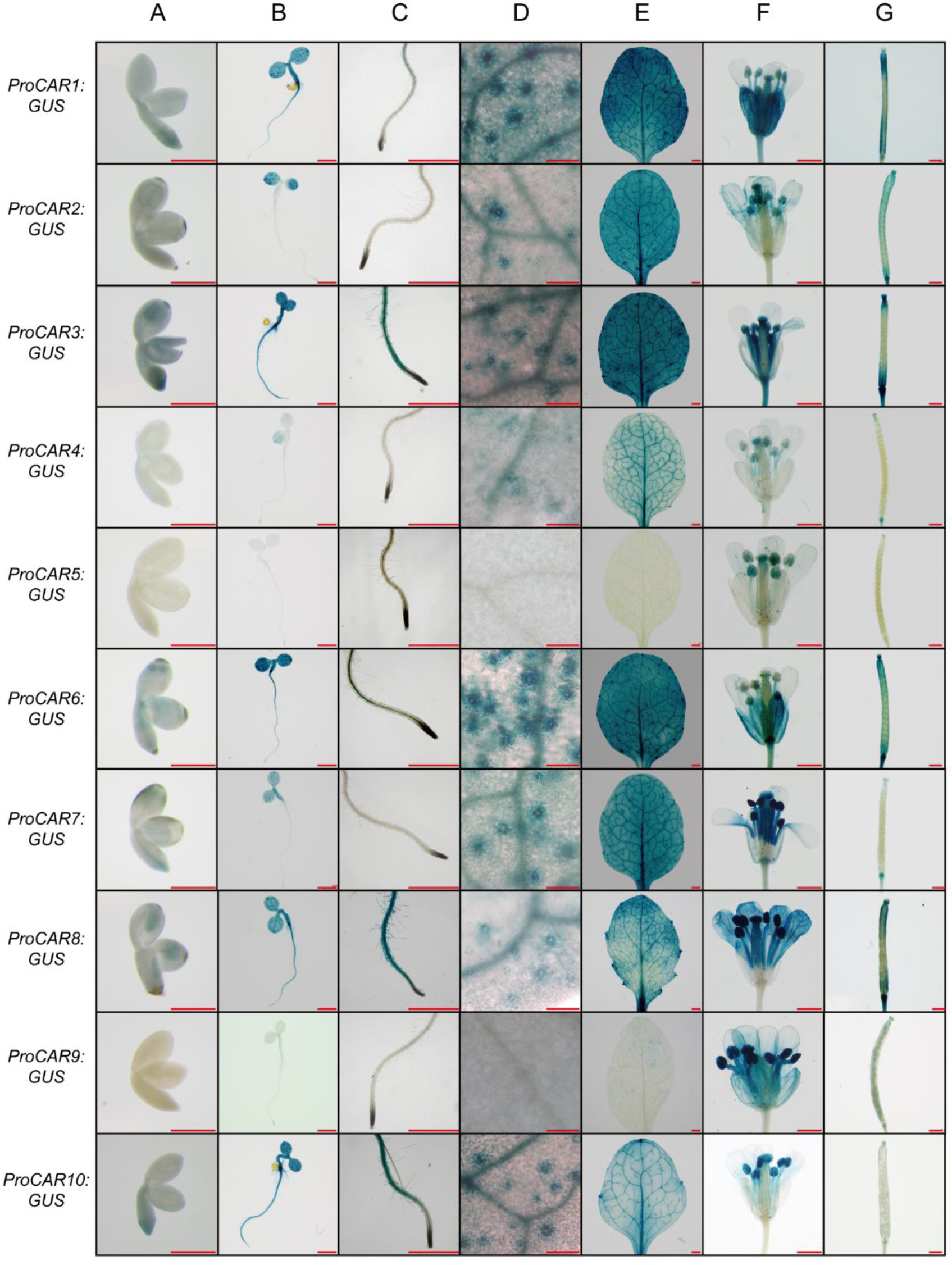
Expression of CAR family genes in different Arabidopsis tissues. Photographs showing GUS expression driven by ProCAR1, ProCAR2, ProCAR3, ProCAR4, ProCAR5, ProCAR6, ProCAR7, ProCAR8, ProCAR9, and ProCAR10 genes in different tissues. (A) Embryos dissected from seeds imbibed for 24 h. (B) 4-d-old seedlings. (C) Primary roots from 4-d-old seedlings. (D) Guard cells in the leaves of 14-d-old seedlings. (E) Leaves of 14-d-old seedlings (F) Flowers. (G) Seed pods. Scale bars, 1 mm (A, B, C, E, F, and G); 0.1 mm (D).

### ABA changes in CAR1-mGFP5 dynamics at the PM

CAR family proteins have been proposed to mediate the recruitment of PYLs for ABA sensing on the membrane. However, direct evidence of its involvement in the ABA signaling response is only available for members of CAR1, CAR4, and CAR9 (Rodriguez et al., 2014; Qin et al., 2019). It is uncertain whether all CAR family members respond to ABA. Therefore, we checked the distribution of CAR proteins affected by ABA in cells, following previous methods (Qin et al., 2019). As shown in Supplemental Figure S1, all CARs-GFP proteins were similarly localized, with fluorescent signals distributed in the nucleus and the perimeter of the cells. In contrast, the distribution of all CAR family members at the PM was increased in the presence of ABA. Thus, considering the remarkable response of CAR1 to ABA (Supplemental Figure S1) and its representation in tissue expression (Figure 1), as well as the interaction with the three core components of ABA signaling (Supplemental Figure S2-S5); moreover, CAR proteins are highly redundant in their response to ABA. Therefore, we chose CAR1 as an example to study the behavior of CAR family proteins in our subsequent studies.

Detecting the kinetic behavior of molecules is critical to understanding their molecular mechanisms. Over the past few decades, the application of single-molecule techniques has allowed the direct detection of heterogeneity between different molecules in living cells at the single-molecule level in real time (Wu et al., 2019; Guo et al., 2021). To investigate the spatial and temporal dynamics of CAR1 in response to ABA at the PM, we generated transgenic *Arabidopsis* expressing a CAR1-mGFP5/CAR1-mCherry fusion protein under the control of the native CAR1 promoter in the *car1* mutant background. Laser scanning confocal microscopy revealed the expression of CAR1-mGFP5/CAR1-mCherry in multiple tissues, including the maturation zone, elongation zone, hypocotyl cells, cotyledon epidermal cells, and guard cells, and it was mainly targeted to the nuclear region and the perimeter of the cell (Supplemental Figure S6A), consistent with previous reports (Rodriguez et al., 2014). The gene expression level and immunoblot analysis confirmed that the selected transgenic lines were standard complementation materials (Supplemental Figures S6B-C).

We employed variable-angle total internal reflection fluorescence microscopy (VA-TIRFM), which produces continuous images with high resolution and a high signal-to-noise ratio to track the dynamics of proteins on the PM (Xing et al., 2019). CAR1-mGFP5 manifested as discrete and diffraction-limited spots with high-speed dynamics (Supplemental Movie S1). To visualize the response of CAR1 to ABA, 4-d-old vertically grown seedlings were transferred to solutions with different ABA concentrations for 10 min. These seedlings were then observed under VA-TIRFM. The single-particle tracking (SPT) technique was used to quantitatively monitor the different dynamic distributions of individual particles in live cells (Wu et al., 2019). As shown in Figures 2A and 2B, the amount of CAR1-mGFP5 on the PM increased with increasing ABA concentrations, consistent with previous reports and Supplemental Figure S1 (Qin et al., 2019). This phenomenon can be explained because the anchoring of CARs to the PM is Ca^2+^ dependent (Rodriguez et al., 2014; Diaz et al., 2016), and ABA can induce an increase in the intracytoplasmic Ca^2+^ concentration.

**Figure 2.**
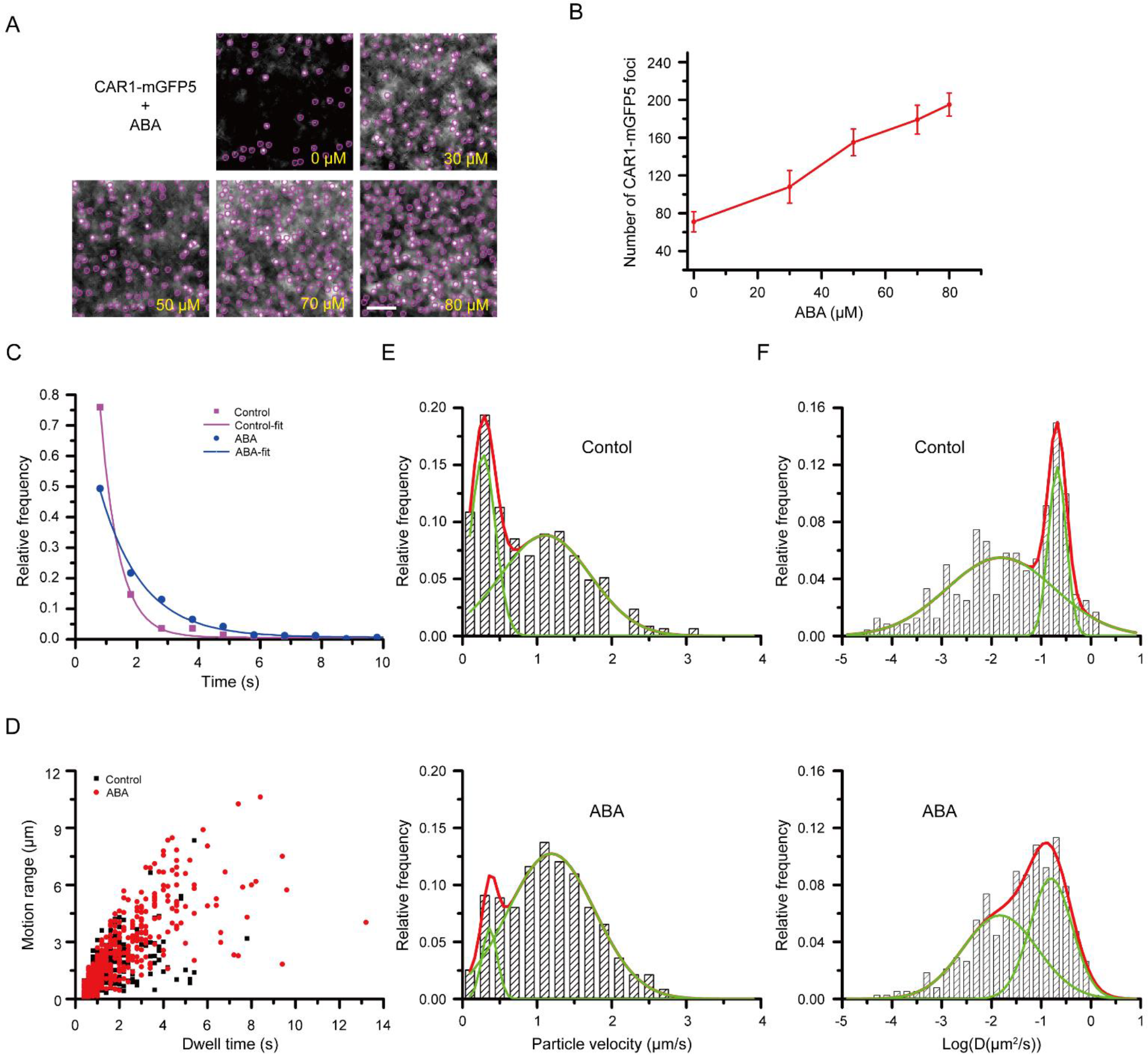
Dynamics of CAR1-mGFP5 on the plasma membrane (PM). (A) Sequential single-particle images for CAR1-mGFP5 at the PM by VA-TIRFM lacking or supplemented with the indicated ABA concentrations. Bars = 3 μm. (B) The number of CAR1-mGFP5 fluorescent spots at the PM in A. Error bars show the SD of three biological replicates. (C) Surface dwell time distribution of CAR1-mGFP5 spots without treatment (n = 705 spots) and with ABA treatment (n = 711 spots). (D) Bubble plots representing CAR1-mGFP5 without treatment (n = 705 spots) or with ABA treatment (n = 711 spots) for motion ranges and dwell times. (E–F) Distribution of particle velocity (E) and diffusion coefficients (F) of CAR1-mGFP5 spots without treatment (n = 681 spots in E and 362 spots in F) or with ABA treatment (n = 711 spots in E and 570 spots in F) at the PM.

Sequential images revealed that individual CAR1-mGFP5 particles lingered on the PM for a brief period of time before quickly disappearing, and in contrast to previous studies on PM proteins, there were a few fluorescent spots that remained and moved on the PM for a long time (Supplemental Movie S1) (Xue et al., 2018). Using SPT analysis, we measured the dwell times of CAR1-mGFP5 under ABA conditions (Supplemental Movie S2). The curves for CAR1-mGFP5 dwell times with or without ABA were fitted to exponential functions, with a τ value of 0.61 s after 10 min of treatment with 0 μM of ABA and a τ value of 1.36 s after 10 min of treatment with 30 μM of ABA (Figure 2C). These data demonstrated that the presence of ABA prolonged the dwell times of CAR1 on the PM. This time period may be needed to complete the response to ABA on the PM.

We used bubble-plot analysis to identify correlations between the motion range and dwell time of CAR1-mGFP5 particles in the presence or absence of ABA. After treatment with 30 μM of ABA for 10 min, the distribution range of CAR1-mGFP5 spots was broader (Figure 2D), indicating that the presence of ABA promoted CAR1 to exhibit longer dwell time and a larger range of motion. We also determined the particle velocity and diffusion coefficients of the CAR1-mGFP5 spots. The particle velocity distribution of untreated CAR1-mGFP5 spots exhibited a bimodal distribution, with 42.9% of spots exhibiting low velocity and 57.1% exhibiting high velocity (Ĝ = 0.277 ± 0.012 μm/s and 1.113 ± 0.077 μm/s, respectively; Figure 2E, above). Under the 30 μM of ABA treatment, 20.4% of CAR1-mGFP5 spots exhibited low velocity, and 79.6% exhibited high velocity (Ĝ =0.363 ± 0.031 μm/s and 1.192 ± 0.244 μm/s, respectively; Figure 2E, below). For the diffusion coefficients, without treatment, the two populations had Ĝ_D_ values of 0.015 ± 0.004 µm^2^/s (53.5%) and 0.211 ± 0.009 µm^2^/s (46.5%) (Figure 2F, below). Under ABA treatment, the Ĝ_D_ values of the two populations were 0.015 ± 0.027 µm^2^/s (35.5%) and 0.157 ± 0.031 µm^2^/s (64.5%) (Figure 2F, below). Overall, under ABA treatment, the number of CAR1-mGFP5 particles on the PM increased, the dwell time was prolonged, the range of motion was expanded, and the response speed and diffusion coefficients were enhanced. This indicates that, under ABA conditions, the CAR protein responds to ABA by comprehensively enhancing its dynamic behavior, similar to the dynamic behavior changes of the BR receptor BRI1 upon activation by the BR analog 24-epibrassinolide (eBL) (Wang et al., 2015).

### CAR proteins can recruit both PP2Cs and SnRK2s to membranes

Although the relationship between CAR1 and CAR4 and some members of the ABA receptor PYLs has been established (Rodriguez et al., 2014), whether other CAR proteins can interact with PYLs is still unknown. In addition, studies on how PYL transmits signals after receiving ABA signals at the PM are still lacking. The ABA signaling core consists of three components: PYLs, PP2Cs, and SnRK2s. Moreover, similar to PYLs, SnRK2 and PP2C family members are either localized in the nucleus and cytoplasm or localized only in the nucleus (Fujita et al., 2009; Antoni et al., 2012). There is a possibility that CARs could bring these two to the membrane. If this is the case, ABA signaling can be accomplished across the membrane, which is economical for the cell.

To comprehensively validate the interaction between CARs and core ABA signaling factors, the bimolecular fluorescence complementation (BiFC) method was used to detect the interactions between CARs-PYLs, CARs-PP2Cs, and CARs-SnRK2s in plant cells (Supplemental Figures S2-S5). Detecting the expression of YFP fluorescence and statistically analyzing its fluorescence intensity showed that most combinations had strong fluorescence; however, the expression levels of YFP fluorescence proteins varied, reflecting the different strengths of the interaction between CARs and core ABA signaling factors. Moreover, in terms of interaction localization, the fluorescence emitted by the interactions of CAR family proteins with AHG1, AHG3, and HAI3 were observed only in the nucleus, while the other interactions were in the nucleus and on the PM (or cytosolic). The BiFC results suggested that CAR proteins interacted with the three core components of ABA signaling.

To further confirm the interaction between CAR1 and PP2Cs, and SnRK2s, we employed Y2H, BiFC, and co-immunoprecipitation (Co-IP) experiments. In the deficient SD medium (-Trp -Leu -His -Ade), colony growth was observed for the combinations of CAR1-AD + HAB1-BD, CAR1-AD + HAB2-BD, and CAR1-AD + SnRK2.2-BD, while the negative controls showed no colony growth (Figure 3A). BiFC analysis revealed that the interaction between CAR1 and ABI2, HAB1, HAI2, SnRK2.2, and SnRK2.3 occurred at the plant PM (Figure 3B). In Co-IP assays, CAR1-mCherry was detected in complexes immunoprecipitated with the anti-GFP antibody from the leaves of transgenic plants co-expressing CAR1-mCherry/HAB1-mGFP5, CAR1-mCherry/HAB2-mGFP5, or CAR1-mCherry/SnRK2.2-mGFP5 (Figure 3C). These data suggest that CAR1 can physically interact with PP2Cs and SnRK2s in plant cells.

**Figure 3.**
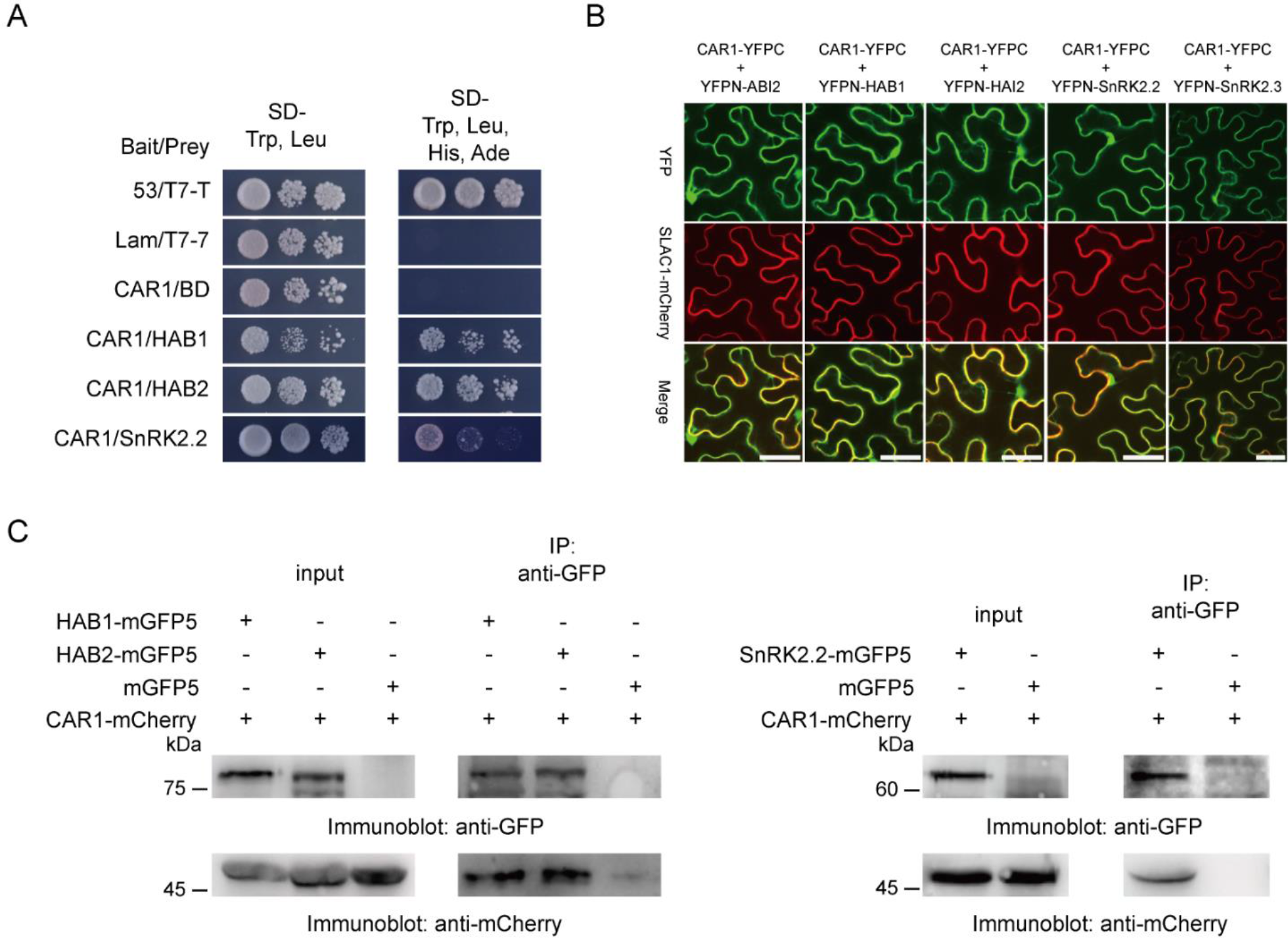
CAR1 interacts with both PP2Cs and SnRK2s. (A) HAB1-BD, HAB2-BD, and SnRK2.2-BD interact with CAR1-AD in Y2H assay. Quadruple selection medium (right) growth is an indication of protein–protein interaction. (B) CAR1 and ABI2, HAB1, HAI2, SnRK2.2, or SnRK2.3 interaction analysis using the BiFC assay. Confocal images of transiently converted Nicotiana benthamiana epidermal cells that co-express CAR1-YFPC with YFPN-ABI2, YFPN-HAB1, YFPN-HAI2, YFPN-SnRK2.2, or YFPN-SnRK2.3 and the PM marker SLAC1-mRuby2 in transiently transformed N. benthamiana epidermal cells. Bars = 50 μm. (C) Co-IP assay confirm the interaction between CAR1 and HAB1, HAB2, and SnRK2.2. Total protein was extracted from 2-week-old plants co-expressing CAR1-mCherry/HAB1-mGFP5, CAR1-mCherry/HAB2-mGFP5, or CAR1-mCherry/SnRK2.2-mGFP5. Plants expressing both CAR1-mCherry and GFP were used as negative controls. An anti-GFP antibody was used to immunoprecipitate the total protein. Blots were examined using anti-GFP or -mCherry probes.

Evidence for the PM recruitment of PYLs by CAR1 has been reported (Rodriguez et al., 2014). Based on the interaction between CAR1 and PP2Cs, and SnRK2s, we wanted to confirm whether CAR1 could recruit PP2Cs and SnRK2s to the PM. SLAC1-mRuby2 was used as a marker of PM localization (Vahisalu et al., 2008). Therefore, we examined the distribution of PP2Cs and SnRK2s proteins at the PM in the presence or absence of CAR1 proteins in the cells as previously described (Rodriguez et al., 2014). First, we co-expressed HAB1-mVenus, HAI2-mVenus, SnRK2.2-mVenus, or SnRK2.3-mVenus with 6Myc as well as SLAC1-mRuby2, and statistically analyzed their co-localization (Figure 4A). As a result, linear Pearson correlation coefficients were calculated to be in the range of 0.60 to 0.73, and non-linear Spearman correlation coefficients were in the range of 0.59 to 0.65 (Figure 4C). We then co-expressed HAB1-mVenus, HAI2-mVenus, SnRK2.2-mVenus, or SnRK2.3-mVenus with CAR1-6Myc (Figure 4B) and found that their co-localization with SLAC1-mRuby2 significantly increased, with linear Pearson correlation coefficients ranging from 0.79 to 0.89 and non-linear Spearman correlation coefficients ranging from 0.79 to 0.88 (Figure 4C). By contrast, the presence of CAR1 protein did not increase the degree of co-localization of mVenus alone with SLAC1-mRuby2. These results suggest that CAR1 may play a role in recruiting PP2Cs and SnRK2s to the PM. To visualize the recruitment of PYL4, HAB1, and SnRK2.2 to the PM by CAR1 at the single practical level, we created stable transgenic *Arabidopsis* lines co-expressing CAR1-mGFP5/PYL4-mCherry, CAR1-mCherry/HAB1-mGFP5, or CAR1-mCherry/SnRK2.2-mGFP5 (Supplemental Figure S7). Dual-color VA-TIRFM showed co-localization and co-diffusion of CAR1-mGFP5 with PYL4-mCherry, CAR1-mCherry with HAB1-mGFP5, and CAR1-mCherry with SnRK2.2-mGFP5 on the PM (Figures 4D–4I). These results indicate that, in addition to ABA receptor PYLs, CAR proteins can recruit two other members of ABA core signaling factors, PP2Cs and SnRK2s, to the cell PM, thereby completing early ABA signaling transduction on the PM.

**Figure 4.**
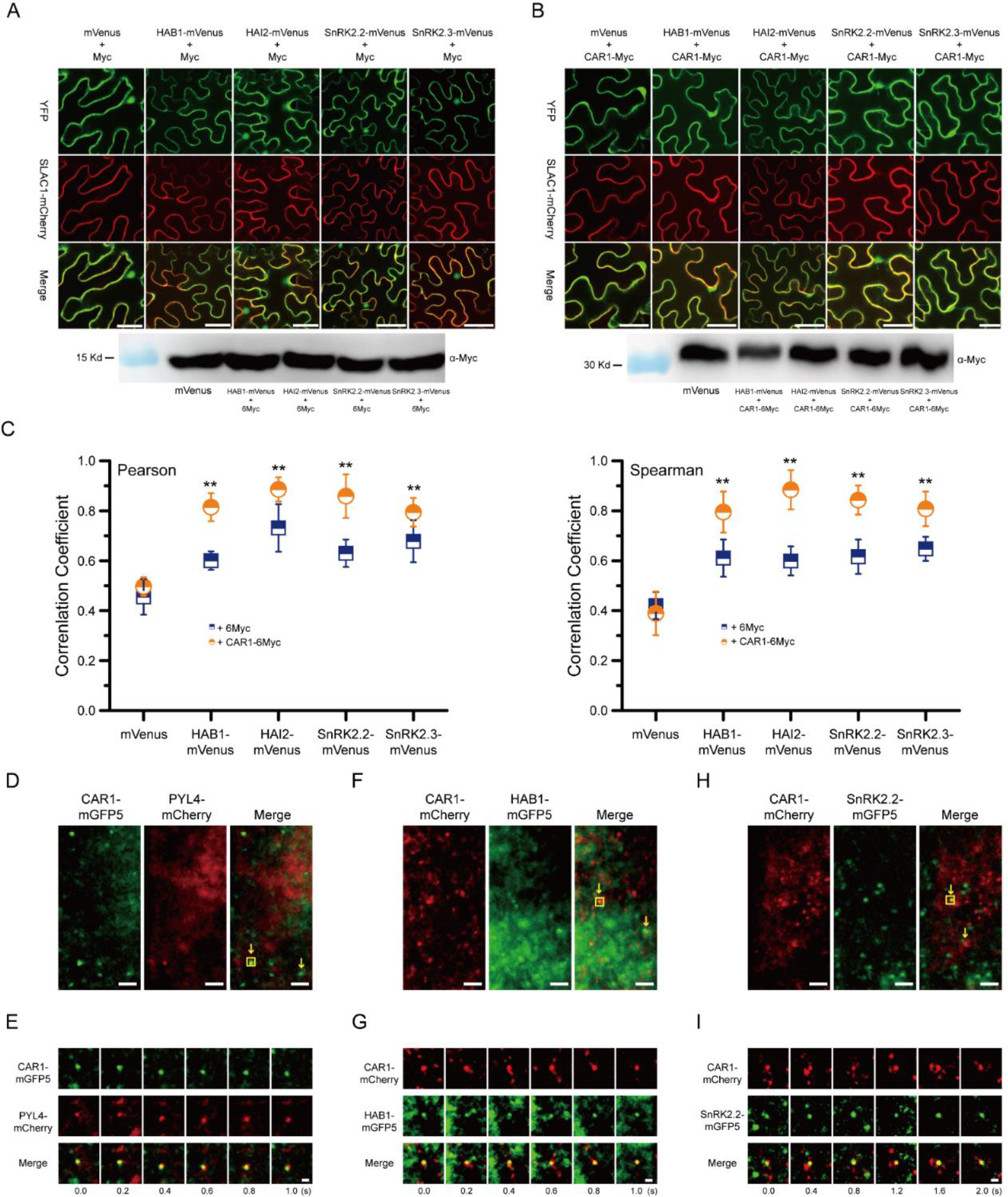
CAR1 recruits the main components of the ABA signaling pathway to membranes. (A) Confocal images showing co-expression of the proteins mVenus, HAB1-mVenus, HAI2-mVenus, SnRK2.2-mVenus, or SnRK2.3-mVenus with 6Myc and the PM marker SLAC1-mRuby2 in transiently transformed Nicotiana benthamiana epidermal cells. Bars = 50 μm. Immunoblot analyses confirm the expression of 6Myc-tags in N. benthamiana epidermal cells (B) Confocal images showing co-expression of the proteins CAR1-6Myc with mVenus, HAB1-mVenus, HAI2-mVenus, SnRK2.2-mVenus, or SnRK2.3-mVenus and the PM marker SLAC1-mRuby2 in transiently transformed Nicotiana benthamiana epidermal cells. Bars = 50 μm. Immunoblot analyses confirm the expression of 6Myc-tagged CAR1 proteins in N. benthamiana epidermal cells. (C) Quantification of the co-localization of mVenus, HAB1-mVenus, HAI2-mVenus, SnRK2.2-mVenus, or SnRK2.3-mVenus with the PM marker SLAC1-mRuby2. Spearman’s rank correlation coefficients or Pearson correlation coefficients were used to calculate co-localization ratios in Image J. Asterisks indicates statistically significant differences, as indicated by the Student’s t-test (**P < 0.01); error bars show the SD of three biological replicates (n ≥ 10). (D) Seedlings expressing CAR1-mGFP5 and PYL4-mCherry were imaged by VA-TIRFM.(E) An example of real-time dynamic observation of CAR1-mGFP5 and PYL4-mCherry co-diffusion (circled puncta in D). (F) Seedlings expressing CAR1-mCherry and HAB1-mGFP5 were imaged by VA-TIRFM. (G) An example of real-time dynamic observation of CAR1-mCherry and HAB1-mGFP5 co-diffusion (circled puncta in F). (H) Seedlings expressing CAR1-mCherry and SnRK2.2-mGFP5 were imaged by VA-TIRFM. (I) An example of real-time dynamic observation of CAR1-mCherry and SnRK2.2-mGFP5 co-diffusion (circled puncta in H). Scale bars represent 2.5 μm (D, F, H); 1 μm (E, G, I).

### *CAR* genes are involved in ABA-mediated seed germination and early seedling development

The above findings indicate the important role of CAR genes in ABA signaling response. Therefore, we hoped to elucidate the role of CARs in ABA-mediated seed germination and early seedling development. To confirm the involvement of CARs in the ABA response, we generated a triple mutant (*car1610*) and two CAR1-overexpressing plants (OE-1 and OE-2). The expression of *CAR1* in these lines was assessed by qRT-PCR (Supplemental Figure S6B). These *Arabidopsis* materials were further tested for germination percentage and cotyledon greening percentage to analyzed their sensitivity to ABA. When grown on MS media, *car1610* and CAR1-overexpressing plants did not obviously differ from the WT plants. However, the germination and cotyledon greenness of the *car1610* mutants were less sensitive to ABA in the presence of ABA, whereas the CAR1-overexpressing plants showed enhanced sensitivity to ABA (Figure 5). In the presence of 0.3 μM ABA, only 25% of WT seeds germinated after 60 h. In contrast, the germination rates of *car1610* seeds were 43%, and CAR1-overexpressing plants were 18%, and 17%, respectively (Figure 5A and 5B). After 10 d of growth in the presence of 0.5 μM ABA, compared with 9% of the WT, the cotyledon greening percentage of the *car1610* was 18%, and CAR1-overexpressing plants lines were 5%, and 4%, respectively (Figure 5C and 5D). Together, these results suggest that *CAR* genes act as positive regulators of ABA signaling during seed germination and early seedling development.

**Figure 5.**
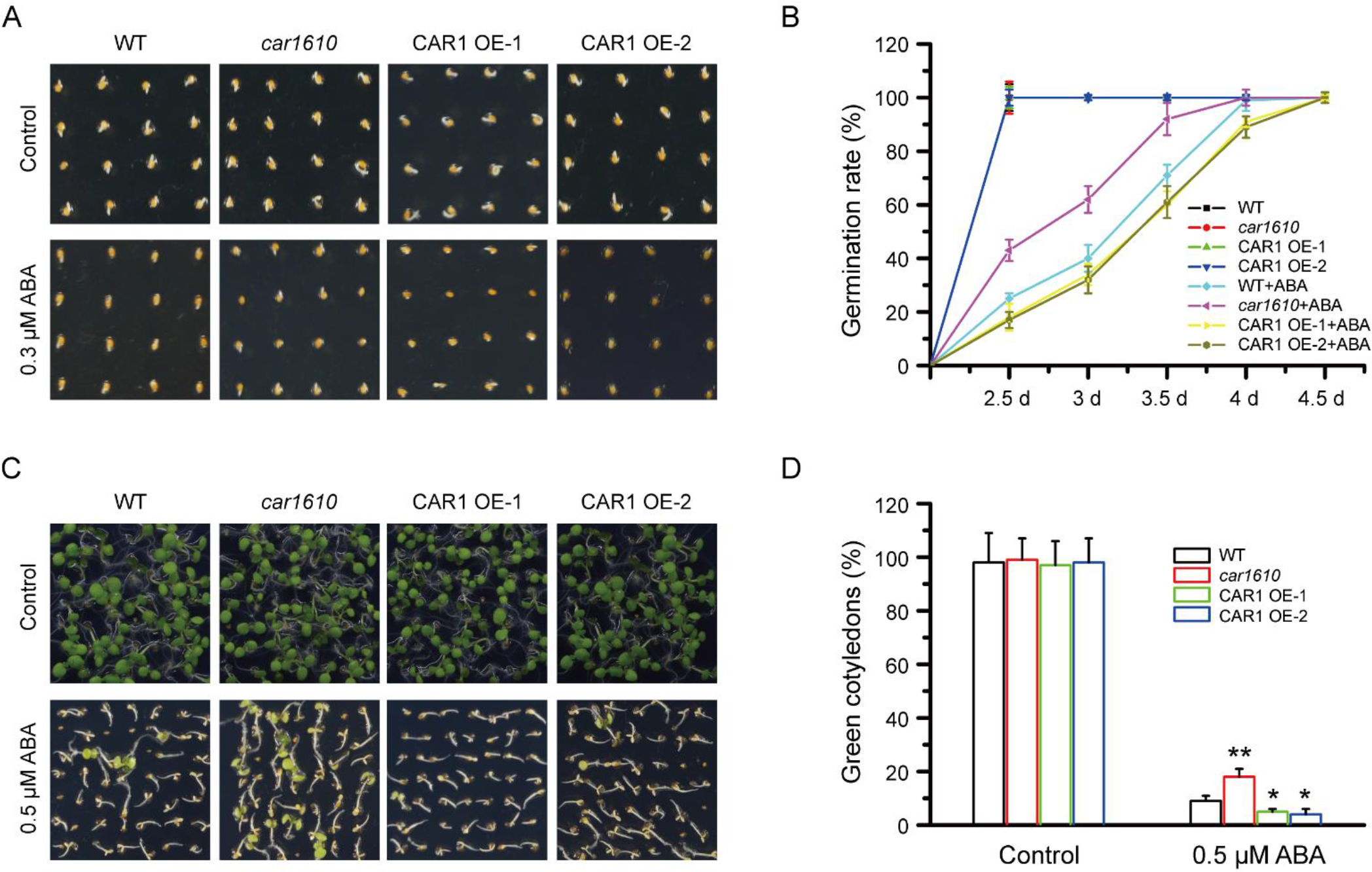
CARs are involved in ABA-mediated seed germination and early seedling development. (A) Representative images of WT, car1610, and two CAR1-overexpressing (OE) lines (#1 and #2) after 60 h of germination on media containing 0 or 0.3 μM ABA. (B) The germination rate of WT, car1610, and two CAR1-OE seeds on media containing 0 or 0.3 μM ABA. Error bars show the SD of three biological replicates. (C) Green cotyledon emergence of the WT, car1610, and two CAR1-OE. Seeds were sown on MS medium without or with 0.5 µM ABA, and photographs were taken 10 d after stratification. (D) Quantification of green cotyledon emergence after 10 d of growth for all genotypes shown in C. Asterisks indicates statistically significant differences, as indicated by the Student’s t-test (*P < 0.05, **P < 0.01); error bars show the SD of three biological replicates.

### Membrane microdomains contribute to CAR1 partitioning

CARs have been confirmed to recruit the main signaling members of the ABA pathway onto the membrane. However, the PM is heterogeneous, making it unclear whether there is a regional distribution of CARs across the PM. PM differs in terms of lipid and protein fractions. Sterol-rich membrane microdomains participate in the polarity of plant cells and tissues, and these domains may regulate the distribution and activity of several membrane proteins (Yu et al., 2020). In addition, these membrane microdomains are essential for signal transduction (Jaillais and Ott, 2020). We employed the sterol-disrupting agent methyl-β-cyclodextrin (MβCD) to examine whether the membrane microdomains were related to the kinetics of CAR1. The number of CAR1-mGFP5 fluorescent spots on the PM was reduced after external MβCD application, and they were reallocated into smaller clusters with greater diameters and a higher fluorescence intensity (Figure 6A-6C). Moreover, the presence of MβCD affected the dynamic response distribution of CAR1 to ABA. After treatment with 30 μM of ABA and MβCD, the CAR1-mGFP5 dwell time curve was fitted as an exponential function with a τ value of 0.69 s (Figure 6D). The particle velocity of CAR1-mGFP5 spots exhibited a bimodal distribution, with 40% of the spots showing low velocity and 60% of the spots showing high velocity (Ĝ = 0.475 ± 0.015 μm/s and 1.657 ± 0.018 μm/s, respectively; Figure 6E). For the diffusion coefficient, the two distributions of Ĝ_D_ values were 0.014 ± 0.007 µm^2^/s (34.5%) and 0.232 ± 0.0212 µm^2^/s (65.5%) (Figure 6F). By comparing the dwell time, particle velocity, and diffusion coefficient of CAR1-mGFP5 under ABA treatment (Figures 3C; 3E, below; 3F, below), we found that the addition of MβCD altered the dynamic behaviors of CAR1 under ABA stress, further supporting that sterol-rich membrane microdomains affect the distribution and dynamic behaviors of CAR1 on the PM and its response to ABA.

**Figure 6.**
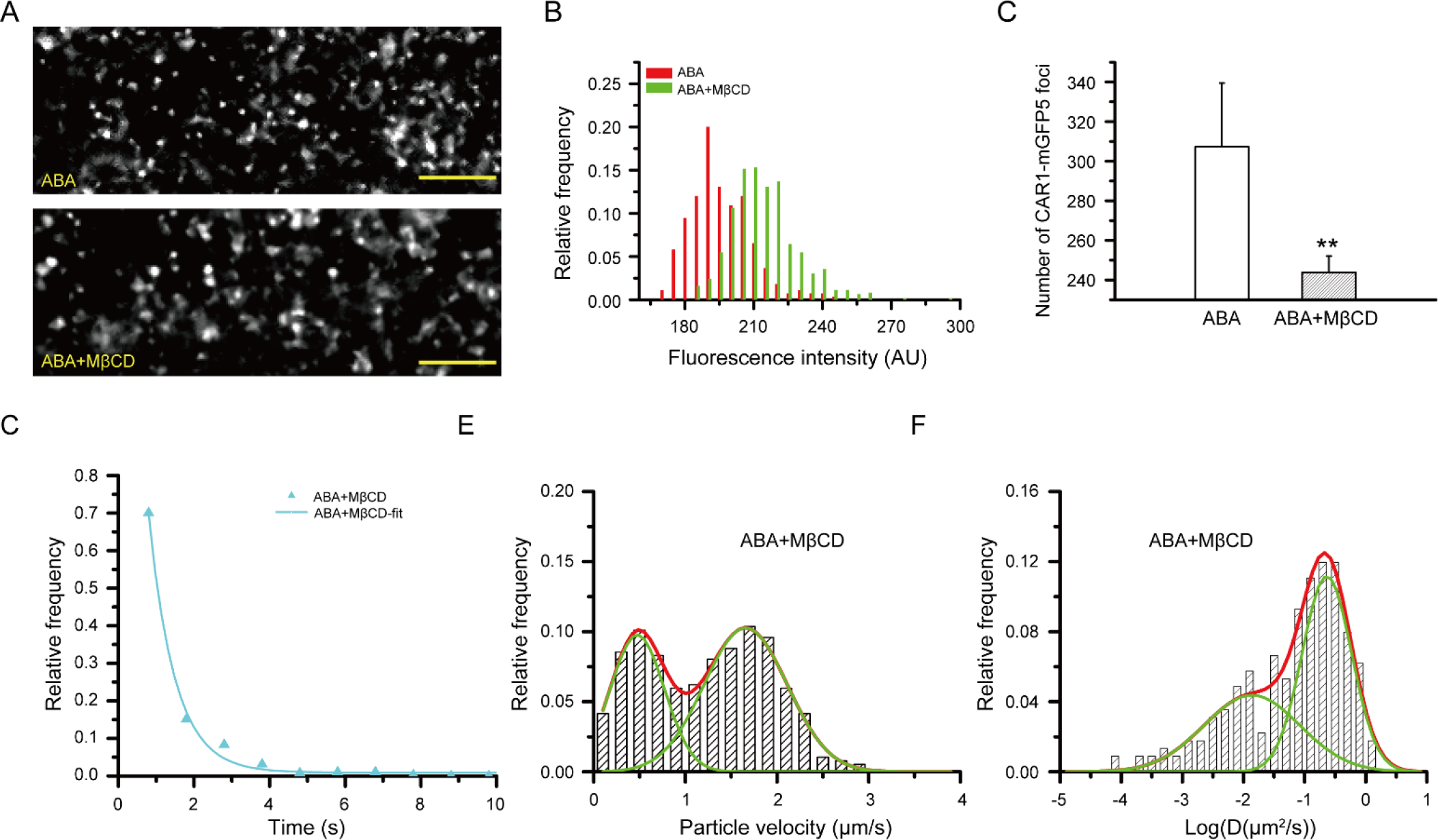
Membrane microdomains contribute to CAR1 localization. (A) VA-TIRFM images of CAR1-mGFP5 in epidermal cells and treated with ABA and both ABA and MβCD. Bars = 5 μm. (B) Distribution of CAR1-mGFP5 spot fluorescence intensities after ABA (n = 621 spots) treatment and both ABA and MβCD (n = 375 spots) treatment. (C) Average fluorescent spot number of CAR1-mGFP5 at the PM after ABA and both ABA and MβCD treatment. Asterisks indicates statistically significant differences, as indicated by the Student’s t-test (**P < 0.01); error bars show the SD of three biological replicates. (D) Surface dwell time distribution of CAR1-mGFP5 spots after MβCD coupled with ABA treatment (n = 576 spots). (E–F) Distribution of particle velocity (E) and diffusion coefficients (F) of CAR1-mGFP5 spots after MβCD coupled with ABA treatment (n = 534 spots in E and 339 spots in F).

By examining the interaction and co-localization of CAR1 with the membrane microdomain marker Flotillin1 (FLOT1), we further investigated the connection between CAR1 and membrane microdomains (Wang et al., 2015). BiFC analysis revealed that the interaction between CAR1 and FLOT1 occurred at the plant PM, consistent with the subcellular localization of FLOT1 (Figure 7A). To confirm the *in vivo* interaction between CAR1 and FLOT1, a Co-IP assay was also applied. The results showed that CAR1-mCherry could be immunoprecipitated with FLOT1-mGFP5. These observations suggest that CAR1 interacts with the FLOT1 protein in plant cells (Figure 7B). Further investigation using dual-color VA-TIRFM imaging of cells co-expressing CAR1-mCherry and FLOT1-mGFP5 revealed some regions with clear fluorescence signals in both GFP and mCherry channels and selected the spots in which CAR1-mCherry co-localized with FLOT1-mGFP5 (Figure 7C). The trajectories of these spots further demonstrated that CAR1-mCherry and FLOT1-mGFP5 co-diffused on the PM for 1.4 s (Figure 7D-7E). These findings provide more credence to the idea that membrane microdomains play a role in CAR1 partitioning.

**Figure 7.**
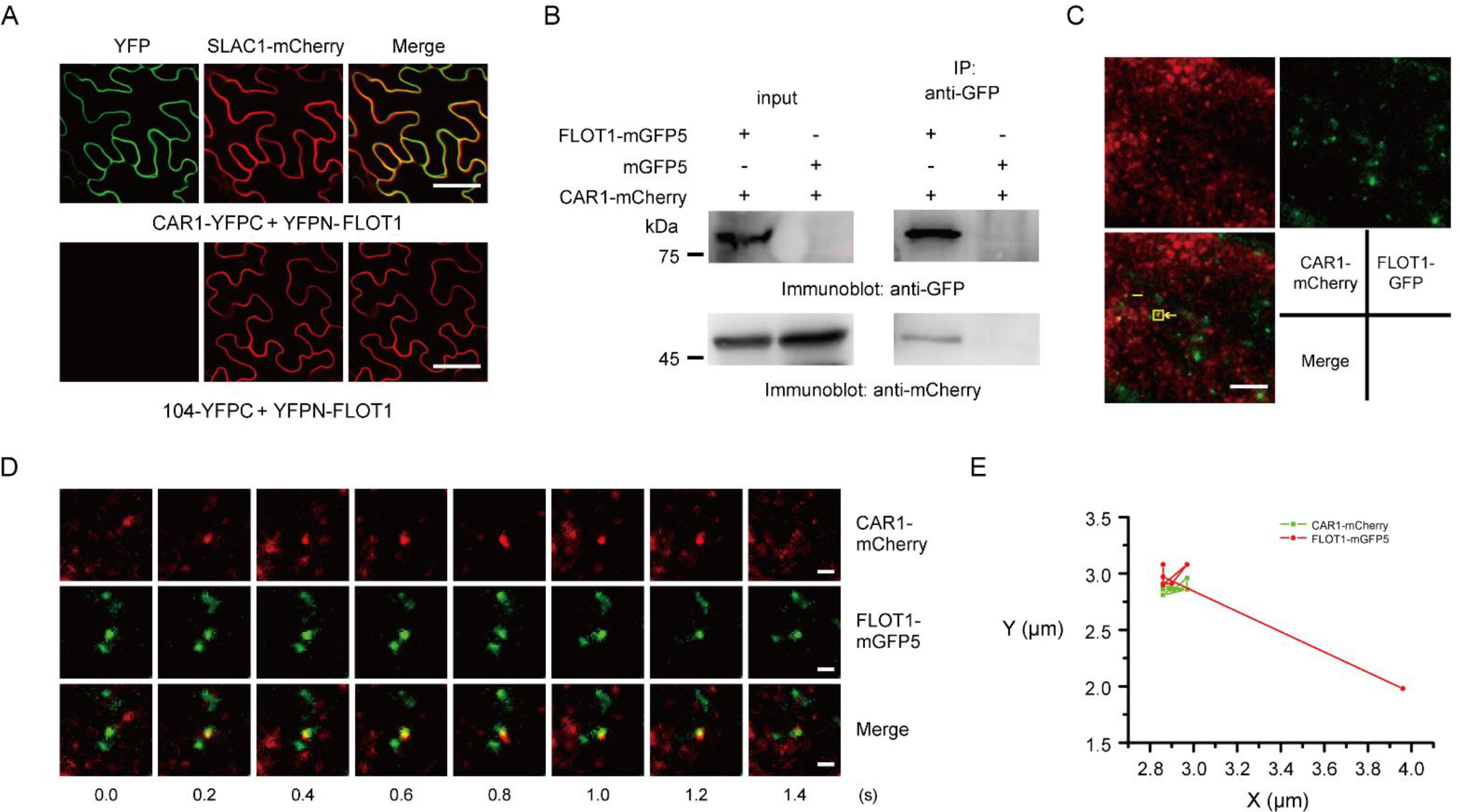
Interaction and co-localization of CAR1-mCherry and FLOT1-mGFP5 foci at the PM. (A) CAR1 and FLOT1 interaction analysis using the BiFC assay. Confocal images of transiently converted Nicotiana benthamiana epidermal cells that co-express CAR1-YFPC, YFPN-FLOT1, and the PM marker SLAC1-mRuby2. Bars = 50 μm. (B) Co-IP assay of the interactions between CAR1 and FLOT1. From transgenic Arabidopsis plants expressing both CAR1-mCherry and FLOT1-mGFP5, total protein was extracted and subjected to immunoprecipitation of the FLOT1 protein with the GFP antibody, followed by immunoblot analysis with the anti-mCherry antibody. (C) VA-TIRFM images of epidermal cells expressing CAR1-mCherry and FLOT1-mGFP5. Yellow arrows show foci that coexist with both GFP and mCherry. Bars = 5 μm. (D) An example of real-time dynamic observation of CAR1-mCherry and FLOT1-mGFP5 co-diffusion (circled puncta in C). Bars = 1 μm. (E) The trajectories of colocalized spots of CAR1-mCherry and FLOT1-mGFP5 in D.

## Discussion

At the PM, many signaling systems rely on information decoding, and membrane-associated proteins and their complexes are essential for controlling this process. How protein complexes are put together and work at membrane locations to alter the identity and dynamics of membrane systems remains a mystery. Peripheral membrane proteins with a C2-domain that can bind calcium and phospholipids can participate in membrane-related signaling by serving as a tether to help protein complexes assemble. CAR proteins, which feature a C2-domain, have recently been identified as important regulators of plant stress responses (Cui et al., 2023). In this work, CAR family members were widely present in *Arabidopsis* and consistently responded to ABA (Figure 1 and Supplemental Figure S1), which implies the potential for powerful functionality and high functional redundancy of the family. However, due to the lack of transmembrane domains in the structure of the CAR proteins, their functioning on the PM is expected to be a relatively brief and rapid process. This means that it can be difficult to understand the dynamics and activation of CAR1 molecules within live cells during ABA responses. In recent years, single-particle analysis has made it possible to identify molecular tracks and recognize specific processes, such as their stoichiometry, heteromeric interactions, and partitioning inside the highly organized plant cell PM in response to various ligand treatments (Zhang et al., 2019). Therefore, we utilized VA-TIRFM technology to perform a high-precision spatiotemporal analysis of CAR1 dynamics in *Arabidopsis* cells. In this study, we found that individual CAR1-mGFP5 particles appeared as isolated fluorescent spots on the PM and demonstrated dramatically different dynamics (Figure 2). Quantifying protein dynamics in live cells is crucial to acquiring a greater understanding of protein interactions and functions and to developing a full picture of membrane proteins in intricate physiological processes, including receptor signaling (Wu et al., 2019). An analysis of CAR1-mGFP5 dwell time, motion range, particle velocity, and diffusion coefficient on the PM revealed shorter dwell times, less long-range movement, and a significantly higher velocity and diffusion coefficient compared to previously reported membrane-localized proteins (Figure 2) (Wang et al., 2015; Xue et al., 2018). Moreover, the above measurements indicate that the spatiotemporal dynamics of individual CAR1-mGFP5 particles on the PM can be directly measured in living cells, which provides a powerful approach to understanding the dynamics and regulatory mechanisms of CAR1 on the PM. In addition, we evaluated the spatial distribution and complex dynamics of CAR1 in response to ABA. Our results showed that CAR1 responded to ABA by enhancing all of its dynamic properties to achieve a faster and more efficient response, which could be crucial for further signal transduction. As a result, our findings shed light on the control of some unique proteins that do not include transmembrane domains but yet function on the PM and delineate the intricate connections between dynamic behaviors and signal sensing.

The first step in ABA signaling is binding to its receptors. These receptors then form a ternary complex with clade A PP2Cs phosphatases upon ABA detection, which causes the phosphatases to become inactive and release the ABA response cascade. Ion transporters and membrane-associated enzymes are involved in these ABA reactions. Rodriguez et al. (2014) identified a C2-domain protein family that interacts with the ABA receptor PYLs and mediates their clustering on the PM in a Ca^2+^-dependent manner. In this work, we found that CAR family members not only interact with ABA receptors but also interact with two other families involved in ABA signaling, namely PP2Cs and SnRK2s, and recruit them to the PM (Figures 3 and 4). These results support the notion that a complete ABA signaling response system can be established directly on the PM through CAR-mediated recruitment, and that activated SnRK2s may directly regulate ion transport proteins and membrane-associated enzymes, enabling a rapid and efficient plant response. This suggests that CAR plays an important role in ABA signaling, which was also verified during germination and early seedling development in *Arabidopsis* (Figure 5).

According to previous reports and the data in this article, CARs are peripheral membrane proteins that functionally cluster and recruit signaling complexes on the membrane (Rodriguez et al., 2014; Diaz et al., 2016). The lateral compartmentalization of the PMs into separate microdomains results in membrane heterogeneity. These sterol- and sphingolipid-rich membrane microdomains are hypothesized to function as signaling hubs. Super-resolution microscopy has been used to identify membrane microdomains in plants, and there is proof that these microdomains are involved in signal transduction (Xue et al., 2018; Yu et al., 2020). In this study, we discovered that after exposure to the sterol-depleting agent MβCD, the size of CAR1-mGFP5 spots grew, and most of the spots were constrained within small particles. From the dwell time, particle velocity, and diffusion rate of CAR1-mGFP5, it was shown that MβCD significantly weakened CAR1’s response to ABA by reducing the number of membrane raft domains (Figure 6). CAR1-mCherry co-diffused with FLOT1-mGFP5 at the PM, further supporting that CAR1 partitioning depends on sterol-based membrane microdomains (Figure 7).

Taken together, tissue localization analysis showed that CAR family members were widely distributed in *Arabidopsis*. Our single-molecule analysis provided unprecedented spatial and temporal resolution for understanding the dynamic regulation of CAR1 on the PM. Our results showed that CAR1 was heterogeneously distributed on the PM of living cells and that the presence of ABA induced higher dynamic activity of CAR1. Furthermore, our findings showed that CAR1 could bring two earlier ABA signal transduction components, SnRK2s and PP2Cs, to the membrane, in addition to ABA receptor PYLs. The membrane microdomains significantly affected their response to ABA. The above results suggest that the various biological activities of CAR1 on the PM cause dynamic membrane responses. Such perturbations in membrane dynamics are often associated with vesicle trafficking and endocytosis events. We speculate that this vesicle trafficking event is related to vesicle-associated membrane protein 724 (VAMP724), which belongs to the SNARE family (Supplemental Figure S8). Understanding this entire process will help elucidate more signaling pathways involving CARs that occur on the PM. CAR family members have been found to participate in regulating PM localization of proteins, such as AGD12, NPH3, IRT1, YchF1, and FER, in response to blue light tropism, gravitropism, iron nutrition, immune, and abiotic stress signaling (Knauer et al., 2011; Cheung et al., 2013; Dummer et al., 2016; Khan et al., 2019; Dummer et al., 2021; Chen et al., 2023). Our preliminary Y2H screen identified many transporter proteins that localize to the PM (Supplemental Figure S9), including those involved in ABA signaling and those that are not. Therefore, it is speculated that CARs may play an important regulatory role in ABA signaling and other signaling pathways that require signal transduction on or near the PM.

## Materials and Methods

### Plant material and growth conditions

*Arabidopsis thaliana* Columbia-0 (Col-0) ecotypes were used as WT. The *car1* (SALK_038968), *car6* (SALK_134720), *car10* (SALK_122977C), and *pyl4* (SAIL_517_C08) mutants were obtained from the *Arabidopsis* Biological Resource Center (ABRC; http://abrc.osu.edu/). The primers used to identify these mutations are listed in Supplemental Table S1. Before being placed on plates containing half-strength Murashige and Skoog (MS) medium, 1% (w/v) sucrose, and 1% (w/v) agar, the seeds were surface-sterilized with 10% (v/v) sodium hypochlorite and 0.1% (v/v) Triton X-100 for 10 min and rinsed five times with sterilized distilled water. Plates were vernalized at 4°C for 48 h and then put in a culture chamber at 22°C with a light regime of 16-hour white light/8-hour dark. The *N*. *benthamiana* plants utilized in the *Agrobacterium*-mediated transient experiments were cultivated in a plant chamber at 22°C, 60% relative humidity, and a photoperiod of 16 h light/8 h dark.

### Plasmid construction and plant transformation

To generate *ProCAR1*, *ProCAR2*, *ProCAR3*, *ProCAR4*, *ProCAR5*, *ProCAR6*, *ProCAR7*, *ProCAR8*, *ProCAR9*, and *ProCAR10:GUS* genes, the upstream region of the ATG sequence that was amplified was roughly 1.5 kb to minimize overlapping with the regulatory regions of nearby genes. The promoter regions were amplified from genomic DNA using PCR and subcloned into the vector *pCAMBIA3301-GUS*. The primer sequences used are listed in Supplemental Table S1. The various constructs containing *ProCAR1-10:GUS* genes were introduced to *Agrobacterium tumefaciens* GV3101 and utilized to transform WT plants using the floral dip method (Clough and Bent, 1998). To identify transgenic plants, transformed plant seeds were collected and plated on basta (10 μg/mL) selection media. T3 progeny homozygous for the selection marker were then employed for further research. As previously mentioned, a GUS assay was carried out (Gonzalez-Guzman et al., 2012).

The expression vectors for *35s:CAR1-mCherry*, *ProCAR1:CAR1-mCherry*, *ProPYL4:PYL4-mCherry*, *ProCAR1:CAR1-mGFP5*, *ProHAB1:HAB1-mGFP5 ProHAB2:HAB2-mGFP5 ProSnRK2*.*2:SnRK2*.*2-mGFP5*, *ProVAMP724:VAMP724-mGFP5*, and *ProFLOT1:FLOT1-mGFP5* plants were constructed as follows. The coding regions for *CAR1*, *PYL4*, *HAB1*, *HAB2*, *SnRK2*.*2*, *VAMP724*, and *FLOT1* were amplified from an *Arabidopsis* seedling cDNA library and subcloned into the binary plant expression vectors *pCAMBIA1300-mCherry*, *pEGAD-mCherry*, and *pCAMBIA1300-mGFP5* under the control of 35S promoter or the native promoter. The primer sequences used are shown in Supplemental Table S1. The Col-0, *car1* mutant, and *pyl4* mutant were transformed with constructs for *35s:CAR1-mCherry*, *ProFLOT1:FLOT1-mGFP5*, *PCAR1:CAR1-mGFP5*/*ProCAR1:CAR1-mCherry*, and *ProPYL4:PYL4-mCherry* using the *Agrobacterium tumefaciens*-mediated floral dip method (Clough and Bent, 1998). For dual-color VA-TIRFM imaging, transgenic *ProCAR1:CAR1-mGFP5* plants were crossed with *ProPYL4:PYL4-mCherry* plants to produce plants co-expressing the proteins CAR1-mGFP5/PYL4-mCherry. Transgenic *ProCAR1:CAR1-mCherry* plants were crossed with *ProFLOT1:FLOT1-mGFP5* plants to produce plants co-expressing the proteins CAR1-mCherry/ FLOT1-mGFP5. Transgenic *ProCAR1:CAR1-mCherry* lines were transformed with the vector containing *ProHAB1:HAB1-mGFP5*, *ProHAB2:HAB2-mGFP5*, *ProSnRK2*.*2:SnRK2*.*2-mGFP5*, and *ProVAMP724:VAMP724-mGFP5* to produce plants co-expressing the proteins CAR1-mCherry/HAB1-mGFP5, CAR1-mCherry/HAB2-mGFP5, CAR1-mCherry/SnRK2.2-mGFP5, and CAR1-mCherry/VAMP724-mGFP5. Transgenic plants were selected on solid medium (0.6% agar), half-strength MS medium containing 25 mg/L of hygromycin for *ProCAR1:CAR1-mGFP5*, *ProHAB1:HAB1-mGFP5*, *ProHAB2:HAB2-mGFP5*, *ProSnRK2*.*2:SnRK2*.*2-mGFP5*, *ProVAMP724:VAMP724-mGFP5*, and *ProFLOT1:FLOT1-mGFP5* and 10 μg/mL of basta for *ProCAR1:CAR1-mCherry* and *ProPYL4:PYL4-mCherry*.

The expression vectors for *35S:CAR1-10-GFP*, *CAR1-6Myc*, *HAB1-mVenus*, *HAI2-mVenus*, *SnRK2*.*2-mVenus*, *SnRK2*.*3-mVenus*, and SLAC1-mRuby2 were constructed as follows. The coding regions for *CAR1-10*, *HAB1*, *HAI2*, *SnRK2*.*2*, *SnRK2*.*3*, and *SLAC1* were amplified from an *Arabidopsis* seedling cDNA library and subcloned into the vectors *pCAMBIA1300-GFP*, *pCAMBIA1307-Myc*, *pCAMBIA1300-mVenus*, and *pCAMBIA1300-mRuby2* under the control of the 35S promoter. The primer sequences used are listed in Supplemental Table S1.

### Split-luciferase complementation assay

The split-luciferase complementation assay was performed as previously described (Lou et al., 2020). The CDS of CAR1 was cloned into the 35S promoter driving pCAMBIA1300-CLuc and that of VAMP724 and cloned into the 35S promoter driving pCAMBIA1300-NLuc. The constructs were transformed into *Agrobacterium* GV3101, co-injected into *N*. *benthamiana* leaves, and cultivated for 48–72 h. The images were captured using a CCD camera (FUSION FX EDGE SPECTRA; VILBER, Deutschland, Frankfurt) after spraying 1 mM of D-luciferin on the leaves. The primer sequences are listed in Supplemental Table S1.

### Y2H assay

The cDNA of *Arabidopsis CAR1* was cloned into pGADT7, and the cDNA of *Arabidopsis HAB1*, *HAB2*, and *SnRK2*.*2* was cloned into pGBDT7. Pair vectors of CAR1-AD and HAB1-BD, CAR1-AD and HAB2-BD, CAR1-AD and SnRK2.2-BD, or CAR1-AD and empty vector pGBDT7 were co-transformed into yeast strain Y2HGold. The yeast cells were grown on a -2 SD (-Trp -Leu) or -4 SD (-Trp -Leu -His -Ade) selective medium for 3 days. Supplemental Table S1 lists the primers used for the Y2H assays.

### BiFC assay

The coding sequence of CAR1-10 was amplified and subcloned as a SacI–KpnI fragment into pXY104-YFPC, and the coding sequences of PYL1-13, PYR1, ABI1, ABI2, AHG1, AHG3, HAB1, HAB2, HAI1, HAI2, SnRK2.2, SnRK2.3, SnRK2.6, FLOT1, and VAMP724 were amplified and subcloned as BamHI–SalI fragments into pXY106-YFPN vector, and the constructs were introduced into *Agrobacterium tumefaciens* GV3101 and then co-transformed into *N*. *benthamiana* leaves. SLAC1-mCherry was expressed in *N*. *benthamiana* leaves as a marker protein for PM localization. Plants were grown and allowed to recover for 2 d. The fluorescence images were captured using a scanning confocal microscope (Andor Dragonfly 200; Andor Technology, Belfast, UK). Supplemental Table S1 lists the primers used for the BiFC assays.

### Co-IP assay

The Co-IP assay followed a previously described method (Nie et al., 2022; Zhan et al., 2023). Briefly, the total proteins for CAR1-mCherry/HAB1-mGFP5, CAR1-mCherry/HAB2-mGFP5, CAR1-mCherry/SnRK2.2-mGFP5, CAR1-mCherry/VAMP724-mGFP5, and CAR1-mCherry/FLOT1-mGFP5 transgenic plants were extracted and incubated with anti-GFP beads (Sigma-Aldrich) for 2 h at 4°C. The samples were washed five times with extraction buffer and then immunoblotted with anti-Cherry antibodies (Sigma-Aldrich).

### Gene expression analysis

TRIzol reagent (Tiangen, China) was used to isolate the total RNA from 10-d-old seedlings. The first-strand cDNA was reverse transcribed with PrimeScript™ RT Reagent Kit with gDNA Eraser (TaKaRa, Japan). Quantitative real-time PCR was run on a Roche LightCycler® 480 II real-time PCR detection system. Supplemental Table S1 lists the primers used for the RT-qPCR assays.

### Germination assays

Arabidopsis seeds were sown on half-strength MS media without or with the indicated concentration of ABA (Sigma-Aldrich), maintained in darkness at 4°C for 2 d, and then transferred to a growth chamber. Seeds with radicle protruding from the seed coat were recorded as germinated. The percentage of green cotyledons was determined at indicated time points (Zhao et al., 2020).

### Drug Treatments

ABA and MβCD were obtained from Sigma-Aldrich. ABA was used from ethanol-dissolved stock solutions. MβCD was prepared in deionized water. For drug treatment, the chemicals were used at the following working concentrations: MβCD, 10 mM; ABA, 10 μM; ABA, 30 μM; ABA, 50 μM; ABA, 70 μM; ABA, 80 μM. All of these chemicals were diluted using H_2_O, and the chemical treatments were performed as reported previously (Wang et al., 2015).

### VA-TIRFM and fluorescence image analysis

An inverted microscope (IX-83; Olympus, Japan) equipped with a 100× oil-immersion objective (numerical aperture = 1.45; Olympus, Japan) and a total internal reflective fluorescence illuminator were used to observe seedlings. GFP or mCherry proteins were excited with 488- or 561-nm laser lines, and two band-pass filters (525/50 and 630/75) were applied to pass through the emission fluorescence signals. For recording, a back-illuminated sCMOS camera (Prime 95B; TELEDYNE PHOTOMETRICS, TUCSON, USA) was used. CellSens Dimension 2 (Olympus, Japan) was used to acquire the images, with a 200-ms exposure time. According to the previously mentioned spatial and temporal global particle assignment, single particle tracking was carried out (Tinevez et al., 2017). The previously described methods were used to analyze surface dwell time, particle velocity, motion range, and diffusion coefficient (Wang et al., 2015; Xue et al., 2018). The fluorescence intensities of single particles were calculated based on previously described methods (Wang et al., 2015).

### Confocal laser scanning microscopy

An Andor Dragonfly 200 laser scanning microscope was used to capture GFP/YFP and mCherry fluorescence images. GFP/YFP was visualized at 488 nm, with detection between 503 and 540 nm; mCherry was visualized at 561 nm and detected between 573 and 615 nm. Images were taken at the same laser intensity to compare the intensities between genotypes. Confocal images were extracted using Fusion image-processing software.

To quantify the co-localization of fluorescent markers on *N*. *benthamiana* leaves co-infiltrated with the described constructs, epifluorescence confocal images of epidermal leaves were combined. The linear Pearson’s correlation coefficient and nonlinear Spearman’s correlation coefficient were determined for fluorescence co-localization (French et al., 2008). For the co-localization analysis, nuclear fluorescent signals were not considered.

To quantify the relative fluorescence intensities of the BiFC experiments, all images were captured using the same laser intensity, pinhole, and gain settings of the confocal microscope to maintain the high reproducibility of the data (laser, 50%; exposure time, 100 ms). Image quantification of relative fluorescence intensities was performed using Fiji software by measuring the fluorescence intensity corrected for mean background fluorescence subtracted from corresponding areas showing no YFP fluorescence.

### GO annotation

WebGestalt (WEB-based gene set analysis tools) was used to undertake Gene Ontology (GO) analyses (Liao et al., 2019).

### Accession numbers

The *Arabidopsis* Information Resource database contains sequence information from this study, with the following accession numbers: The *Arabidopsis* Information Resource database contains sequence information from this study with the following accession numbers: *CAR1* to *CAR10* (At5g37740, At1g66360, At1g73580, At3g17980, At1g48590, At1g70800, At1g70810, At1g23140, At1g70790, At2g01540); *PYL1* to *PYL13* (At5g46790, At2g26040, At1g73000, At2g38310, At5g05440, At2g40330, At4g01026, At5g53160, At1g01360, At4g27920, At5g45860, At5g45870, At4g18620); *PYR1* (At4g17870); *ABI1*, AT4G26080; *ABI2*, AT5G57050; *AHG1*, AT5G51760; *AHG3*, AT3G11410; *HAB1*, AT1G72770; *HAB2*, AT1G17550; *HAI2*, AT1G07430; *HAI3*, AT2G29380; *SnRK2.2*, AT3G50500; *SnRK2.3*, AT5G66880; *SnRK2.6*, AT4G33950.

## Supplemental Data

**Supplemental Table S1.** All primer sequences used in the study.

**Supplemental Figure S1.** Co-localization analysis of CARs-GFP and PM marker SLAC1-mRuby2 under ABA treatment.

**Supplemental Figure S2.** BiFC assay of the interactions between CARs and PYLs in *N. benthamiana* leaves.

**Supplemental Figure S3.** BiFC assay of the interactions between CARs and PP2Cs in *N. benthamiana* leaves.

**Supplemental Figure S4.** BiFC assay of the interactions between CARs and SnRK2s in *N. benthamiana* leaves.

**Supplemental Figure S5.** 104-YFPC co-expressed with NYFP-PYLs, NYFP-PP2Cs, or NYFP-SnRK2s in *N. benthamiana* leaves.

**Supplemental Figure S6.** Confocal microscopy images and gene and protein expression of CAR1-related transgenic materials in *Arabidopsis* seedlings.

**Supplemental Figure S7.** Confocal microscopy images of transgenic *Arabidopsis* plants co-expressing CAR1-mGFP5/PYL4-mCherry, CAR1-mCherry/HAB1-mGFP5, or CAR1-mCherry/SnRK2.2-mGFP5.

**Supplemental Figure S8.** CAR1 interacts and co-localizes with VAMP724.

**Supplemental Figure S9.** GO annotation enrichment analysis of CAR1 potential interacting proteins.

**Supplemental Movie S1.** The Movie of Dynamics of CAR1-mGFP5 Foci under Normal Condition.

**Supplemental Movie S2.** The Movie of Dynamics of CAR1-mGFP5 Foci Treated with 30 μM ABA.

**Supplemental Movie S3.** The Movie of Dynamics of CAR1-mGFP5 Foci Treated with ABA and MβCD.

## Acknowledgments

We thank Prof. Jingjing Xing, Prof. Wencheng Liu and Prof. Yuan Zheng of Henan University for their advices and careful review of our manuscript. We would also like to thank Gan Zhang of Henan University for his help in data collection and analysis, and Pan Wang of Xinxiang Medical College for her help in graphing with the figure drawing.

## Author contributions

A.-Y.G., W.-Q.W., and C.-P.S. planned the experiments and wrote the manuscript. A.-Y.G., D.B., Y.L., and J.X. performed the experiments and analyzed the data. W.-Q.W. and C.-P.S. contributed to the experimental plan and data interpretation. A.-Y.G. and W.-Q.W. contributed equally. S.G. and C.-P.S. conceived the project under study, obtained funding and necessary resources, interpreted data sets, and supervised the entire study. All authors have read and approved the final manuscript.

## REFERENCES

Antoni R, Gonzalez-Guzman M, Rodriguez L, Rodrigues A, Pizzio GA, Rodriguez PL (2012) Selective inhibition of clade A phosphatases type 2C by PYR/PYL/RCAR abscisic acid receptors. Plant Physiol 158: 970–980.

Brandt B, Brodsky DE, Xue S, Negi J, Iba K, Kangasjärvi J, Ghassemian M, Stephan AB, Hu H, Schroeder JI (2012) Reconstitution of abscisic acid activation of SLAC1 anion channel by CPK6 and OST1 kinases and branched ABI1 PP2C phosphatase action. PNAS 109: 10593–10598.

Chérel I, Michard E, Platet N, Mouline K, Alcon C, Sentenac H, Thibaud JB (2002) Physical and functional interaction of the *Arabidopsis* K(+) channel AKT2 and phosphatase AtPP2CA. Plant Cell 14: 1133–1146.

Chen K, Li GJ, Bressan RA, Song CP, Zhu JK, Zhao Y (2020) Abscisic acid dynamics, signaling, and functions in plants. J Integr Plant Biol 62: 25–54.

Chen W, Zhou H, Xu F, Yu M, Coego A, Rodriguez L, Lu Y, Xie Q, Fu Q, Chen J, Xu G, Wu D, Li X, Li X, Jaillais Y, Rodriguez P, Zhu S, Yu F (2023) CAR modulates plasma membrane nano-organization and immune signaling downstream of RALF1-FERONIA signaling pathway. New Phytol 237: 2148–2162.

Cheung MY, Li MW, Yung YL, Wen CQ, Lam HM (2013) The unconventional P-loop NTPase OsYchF1 and its regulator OsGAP1 play opposite roles in salinity stress tolerance. Plant Cell Environ 36: 2008–2020.

Cheung MY, Ngo JC, Chen Z, Jia Q, Li T, Gou Y, Wang Y, Lam HM (2020) A structure model explaining the binding between a ubiquitous unconventional G-protein (OsYchF1) and a plant-specific C2-domain protein (OsGAP1) from rice. Biochem J 477: 3935–3949.

Clough SJ, Bent AF (1998) Floral dip: a simplified method for Agrobacterium-mediated transformation of *Arabidopsis thaliana*. Plant J 16: 735–743.

Cui M, Gupta SK, Bauer P (2023) Role of the plant-specific calcium-binding C2-DOMAIN ABSCISIC ACID-RELATED (CAR) protein family in environmental signaling. Eur J Cell Biol 102: 151322.

Diaz M, Sanchez-Barrena MJ, Gonzalez-Rubio JM, Rodriguez L, Fernandez D, Antoni R, Yunta C, Belda-Palazon B, Gonzalez-Guzman M, Peirats-Llobet M, et al. (2016). Calcium-dependent oligomerization of CAR proteins at cell membrane modulates ABA signaling. PNAS 113: E396–405.

Dummer M, Michalski C, Essen LO, Rath M, Galland P, Forreiter C (2016) EHB1 and AGD12, two calcium-dependent proteins affect gravitropism antagonistically in *Arabidopsis thaliana*. J Plant Physiol 206: 114–124.

Dummer M, Spasic SZ, Feil M, Michalski C, Forreiter C, Galland P (2021) Tangent algorithm for photogravitropic balance in plants and Phycomyces blakesleeanus: Roles for EHB1 and NPH3 of *Arabidopsis thaliana*. J Plant Physiol 260: 153396.

French AP, Mills S, Swarup R, Bennett MJ, Pridmore TP (2008) Colocalization of fluorescent markers in confocal microscope images of plant cells. Nat Protoc 3: 619–628.

Fujita Y, Nakashima K, Yoshida T, Katagiri T, Kidokoro S, Kanamori N, Umezawa T, Fujita M, Maruyama K, Ishiyama K, et al. (2009) Three SnRK2 protein kinases are the main positive regulators of abscisic acid signaling in response to water stress in *Arabidopsis*. Plant Cell Physiol 50: 2123–2132.

Geiger D, Maierhofer T, Al-Rasheid KA, Scherzer S, Mumm P, Liese A, Ache P, Wellmann C, Marten I, Grill E, et al. (2011) Stomatal closure by fast abscisic acid signaling is mediated by the guard cell anion channel SLAH3 and the receptor RCAR1. Sci Signal 4: ra32.

Gonzalez-Guzman M, Pizzio GA, Antoni R, Vera-Sirera F, Merilo E, Bassel GW, Fernandez MA, Holdsworth MJ, Perez-Amador MA, Kollist H, et al. (2012) *Arabidopsis* PYR/PYL/RCAR receptors play a major role in quantitative regulation of stomatal aperture and transcriptional response to abscisic acid. Plant Cell 24: 2483–2496.

Grondin A, Rodrigues O, Verdoucq L, Merlot S, Leonhardt N (2015) Aquaporins contribute to ABA-triggered stomatal closure through OST1-mediated phosphorylation. Plant cell 27: 1945–1954.

Guo AY, Zhang YM, Wang L, Bai D, Xu YP, Wu WQ (2021) Single-molecule imaging in living plant cells: a methodological review. Int J Mol Sci 22: 5071.

Jaillais Y, Ott T (2020) The nanoscale organization of the plasma membrane and its importance in signaling: a proteolipid perspective. Plant Physiol 182: 1682–1696.

Khan I, Gratz R, Denezhkin P, Schott-Verdugo SN, Angrand K, Genders L, Basgaran RM, Fink-Straube C, Brumbarova T, Gohlke H, Bauer P, et al. (2019) Calcium-promoted interaction between the C2-domain protein EHB1 and metal transporter IRT1 inhibits *Arabidopsis* iron acquisition. Plant Physiol 180: 1564–1581.

Knauer T, Dummer M, Landgraf F, Forreiter C (2011) A negative effector of blue light-induced and gravitropic bending in *Arabidopsis*. Plant Physiol 156: 439–447.

Komatsu K, Suzuki N, Kuwamura M, Nishikawa Y, Nakatani M, Ohtawa H, Takezawa D, Seki M, Tanaka M, Taji, T, et al. (2013) Group A PP2Cs evolved in land plants as key regulators of intrinsic desiccation tolerance. Nat Commun 4: 2219.

Liao Y, Wang J, Jaehnig EJ, Shi Z, Zhang B (2019) WebGestalt 2019: Gene set analysis toolkit with revamped UIs and APIs. Nucleic Acids Res 47: W199–W205.

Lou L, Yu F, Tian M, Liu G, Wu Y, Wu Y, Xia R, Pardo JM, Guo Y, Xie Q (2020) ESCRT-I component VPS23A sustains salt tolerance by strengthening the SOS module in *Arabidopsis*. Mol Plant 13: 1134–1148.

Michalski C, Dümmer M, Galland P, Forreiter C (2017) Impact of EHB1 and AGD12 on root and hypocotyl phototropism in *Arabidopsis thaliana*. J Plant Growth Regul 36: 660–668.

Nguyen QTC, Lee SJ, Choi SW, Na YJ, Song MR, Hoang QTN, Sim SY, Kim MS, Kim JI, Soh MS, et al. (2019) *Arabidopsis* RAF-like kinase Raf10 is a regulatory component of core ABA signaling. Mol Cells 42: 646–660.

Nie K, Zhao H, Wang X, Niu Y, Zhou H, Zheng Y (2022) The MIEL1-ABI5/MYB30 regulatory module fine tunes abscisic acid signaling during seed germination. J Integr Plant Biol 64: 930–941.

Qin T, Tian Q, Wang G, Xiong L (2019) LOWER TEMPERATURE 1 enhances ABA responses and plant drought tolerance by modulating the stability and localization of C2-domain ABA-related proteins in *Arabidopsis*. Mol plant 12: 1243–1258.

Rodriguez L, Gonzalez-Guzman M, Diaz M, Rodrigues A, Izquierdo-Garcia AC, Peirats-Llobet M, Fernandez MA, Antoni R, Fernandez D, Marquez JA, et al. (2014) C2-domain abscisic acid-related proteins mediate the interaction of PYR/PYL/RCAR abscisic acid receptors with the plasma membrane and regulate abscisic acid sensitivity in *Arabidopsis*. Plant cell 26: 4802–4820.

Sirichandra C, Gu D, Hu HC, Davanture M, Lee S, Djaoui M, Valot B, Zivy M, Leung J, Merlot S, et al. (2009) Phosphorylation of the *Arabidopsis* AtrbohF NADPH oxidase by OST1 protein kinase. FEBS Lett 583: 2982–2986.

Tinevez JY, Perry N, Schindelin J, Hoopes GM, Reynolds GD, Laplantine E, Bednarek SY, Shorte SL, Eliceiri KW (2017) TrackMate: An open and extensible platform for single-particle tracking. Methods 115: 80–90.

Tischer SV, Wunschel C, Papacek M, Kleigrewe K, Hofmann T, Christmann A, Grill E (2017) Combinatorial interaction network of abscisic acid receptors and coreceptors from *Arabidopsis thaliana*. PNAS 114: 10280–10285.

Vahisalu T, Kollist H, Wang YF, Nishimura N, Chan WY, Valerio G, Lamminmäki A, Brosché M, Moldau H, Desikan R, et al. (2008) SLAC1 is required for plant guard cell S-type anion channel function in stomatal signaling. Nature 452: 487–491

Wang L, Li H, Lv X, Chen T, Li R, Xue Y, Jiang J, Jin B, Baluska F, Samaj J, et al. (2015) Spatiotemporal dynamics of the BRI1 receptor and its regulation by membrane microdomains in living *Arabidopsis* cells. Mol Plant 8: 1334–1349.

Wang P, Xue L, Batelli G, Lee S, Hou YJ, Van Oosten MJ, Zhang H, Tao WA, Zhu JK (2013) Quantitative phosphoproteomics identifies SnRK2 protein kinase substrates and reveals the effectors of abscisic acid action. PNAS 110: 11205–11210.

Winter D, Vinegar B, Nahal H, Ammar R, Wilson GV, Provart NJ (2007) An “Electronic Fluorescent Pictograph” browser for exploring and analyzing large-scale biological data sets. PloS one 2: e718.

Wu WQ, Zhu X, Song, CP (2019) Single-molecule technique: a revolutionary approach to exploring fundamental questions in plant science. New Phytol 223: 508–510.

Xing J, Li X, Wang X, Lv X, Wang L, Zhang L, Zhu Y, Shen Q, Baluska F, Samaj J, et al. (2019) Secretion of phospholipase Dδ functions as a regulatory mechanism in plant innate immunity. Plant cell 31, 3015–3032.

Xue Y, Xing J, Wan Y, Lv X, Fan L, Zhang Y, Song K, Wang L, Wang X, Deng X, et al. (2018) *Arabidopsis* blue light receptor phototropin 1 undergoes blue light-induced activation in membrane microdomains. Mol Plant 11: 846–859.

Yokotani N, Ichikawa T, Kondou Y, Maeda S, Iwabuchi M, Mori M, Hirochika H, Matsui M, Oda K (2009) Overexpression of a rice gene encoding a small C2 domain protein OsSMCP1 increases tolerance to abiotic and biotic stresses in transgenic *Arabidopsis*. Plant Mol Biol 71: 391–402.

You C, Li C, Ma M, Tang W, Kou M, Yan H, Song W, Gao R, Wang X, Zhang Y, et al. (2022) A C2-domain abscisic acid-related gene, IbCAR1, positively enhances salt tolerance in Sweet Potato (Ipomoea batatas (L.) Lam.). Int J Mol Sci 23: 9680.

Yu M, Cui Y, Zhang X, Li R, Lin J (2020). Organization and dynamics of functional plant membrane microdomains. Cell Mol Life Sci 77: 275–287.

Zhan Q, Shen J, Nie K, Zheng Y (2023) MIW1 participates in ABA signaling through the regulation of MYB30 in *Arabidopsis*. Plant Sci 332: 111717.

Zhang X, Cui Y, Yu M, Su B, Gong W, Baluska F, Komis G, Samaj J, Shan X, Lin J (2019) Phosphorylation-mediated dynamics of nitrate transceptor NRT1.1 regulate auxin flux and nitrate signaling in lateral root growth. Plant Physiol 181: 480–498.

Zhao H, Nie K, Zhou H, Yan X, Zhan Q, Zheng Y, Song CP (2020) ABI5 modulates seed germination via feedback regulation of the expression of the PYR/PYL/RCAR ABA receptor genes. New Phytol 228: 596–608.

Zhu JK (2016) Abiotic stress signaling and responses in plants. Cell 167: 313–324.

